# Functional evaluation of a natural AAV capsid liver targeting motif in human hepatocytes

**DOI:** 10.64898/2026.08.20.745184

**Authors:** Carmen Unzu, Amanda X. Chen, Liliana Mancio-Silva, Eric Zinn, Yanhe Wen, Carla Llinares, Andrea Llanos, Cindy Zhu, Allegra Fieldsend, Julio Sanmiguel, Beatrice Bissig-Choisat, Karl-Dimiter Bissig, Ian Alexander, Sangeeta N. Bhatia, Luk H. Vandenberghe

**Affiliations:** Grousbeck Gene Therapy Center, Schepens Eye Research Institute and Massachusetts Eye and Ear Infirmary, Boston, MA, USA: Ocular Genomics Institute, Department of Ophthalmology, Harvard Medical School, Boston, MA, USA; DNA & RNA Medicine Division, CIMA, Universidad de Navarra. Pamplona, Spain; Department of Biological Engineering, MIT, Cambridge, MA, USA; David H. Koch Institute for Integrative Cancer Research, MIT, Cambridge, MA, USA; Institute for Medical Engineering and Science, Cambridge, MIT, MA, USA; Translational Vectorology Group, Children’s Medical Research Institute, University of Sydney, Sydney, NSW, Australia; Duke University Medical Center, Durham, NC, USA; Marble Center for Cancer Nanomedicine, Massachusetts Institute of Technology, Cambridge, MA 02139, USA; Wyss Institute for Biologically Inspired Engineering, Harvard University, Boston, MA, USA; Department of Electrical Engineering and Computer Science, MIT, Cambridge, USA; Howard Hughes Medical Institute, Chevy Chase, MD, USA; Ludwig Center at MIT’s Koch Institute for Integrative Cancer Research, Massachusetts Institute of Technology, Cambridge, MA 02139, USA; Broad Institute of MIT and Harvard, Cambridge, MA 02142, USA

**Keywords:** liver, gene therapy, adeno-associated viral vectors, AAV, human models, liver detargeting

## Abstract

**Background&Aims:** Adeno-associated virus (AAV) vectors are attractive delivery vehicles for therapeutic gene delivery, and a notable feature of most AAVs is their natural tropism for the liver, which leads to significant hepatic uptake following systemic administration. In previous work, we identified 266G as a conserved motif on a variable region on the capsid of many commonly used AAV variants that controls liver uptake in both mice and non-human primates. This single amino acid could be functionally leveraged to engineer AAVs to either de-target from or enhance tropism to the liver. Here, we explored whether these observations extended to the human context.

**Methods:** Two human hepatocyte models were tested: Fah^−/−^/Rag2^−/−^/Il2rg^−/−^ (FRG) mice with humanized livers and a bioengineered human microliver platform *in vitro*. A barcoded AAV capsid library including standard control serotypes were used to assess the role of the 266G motif on gene transfer and transgene expression in both liver systems.

**Results:** *In vivo*, 266G containing AAVs indeed targeted human hepatocytes superiorly, with some noted dependency on the degree of human-hepatocyte replacement in the chimeric mouse model. Initial studies in the micropatterned primary human hepatocyte co-culture model however demonstrated enrichment of heparin-binding AAVs, and not 266G variants. Notably, incorporation of polyethylene glycol (PEG) into the system modified the AAV transduction potential of those capsids including the liver-targeting motif, recapitulating the hepatocyte transduction pattern observed *in vivo*. Importantly, when PEG was used, the two human models, both at the DNA and RNA level, did correlate significantly.

**Conclusions:** Our results showed the potential of a combinatorial AAV library for model validation and revealed the human microliver platform-PEG as a reliable system for the development of AAV therapeutics.

## INTRODUCTION

Recombinant adeno-associated viral vectors (AAV) are extensively used for *in vivo* delivery of nucleic acids in clinical gene therapy. Most AAV serotypes are naturally hepatotropic^1^, which makes them the preferred viral vector for liver-directed gene medicine. Nevertheless, developing AAV-delivered therapeutics for human liver disease is still limited by the high doses that are sometimes required,^2^ liver toxicity associated with them^2,3^, as well as immune system activation^3^. Conversely, for tissue targets in gene therapy outside of the liver, the need to bypass this organ is desired to avoid liver-associated toxicities and minimize off-target attrition of the input vector. Engineering AAV capsids following strategies from rational to artificial intelligence (AI)-driven design has become an important pursuit to identify clinically desirable vectors that overcome these translational limitations^4,5^.

The major bottleneck in establishing an efficient AAV drug discovery pipeline is the lack of reproducible preclinical systems that faithfully predict capsid performance in the human liver. Historically, animal models have been the preferred choice for studying AAV fitness and biology and, for the development of liver-directed gene therapies, mouse models with the possibility of liver humanization, such as the Fah^−/−^/Rag2^−/−^/Il2rg^−/−^ (FRG) mice^6^ or the Il2rg−/−/Rag2−/−/Fah−/−/Aavr−/− (TIRFA) mice^7^, have been studied extensively for AAV capsid discovery^7,8^. Although these mice present additional challenges including variability in the liver replacement index and lack of a competent immune system^6-8^.

Importantly, with the increasing number of AAV libraries generated to identify liver-directed or liver-detargeted variants^9-12^, there is a need for high throughput *in vitro* screening that yields for robust and reproducible results. Furthermore, the availability of a reliable *ex vivo* or *in vitro* model for gene therapy applications would reduce unnecessary animal experimentation. Recently, the use of normothermic machine-perfused human livers discarded for transplantation has been proposed as one of the most physiologically relevant *ex-vivo* systems for AAV liver targeting validation^13^. Nevertheless, access to perfused human livers is rare and expensive, experimental success depends on the conditions of the liver upon receipt, and their immune system is still incomplete. As a high throughput *in vitro* alternative, 3D liver organoids, livers-on-chips, and micro-livers have arisen as a scalable option for AAV screening and mechanistic studies^14,15^. Although these systems come with their own limitations such as missing a full vascular system or an immature hepatocyte phenotype, which may misrepresent hepatocyte polarity, receptor density, extracellular matrix therefore affecting natural AAV access routes^16,17^. In fact, AAV cell transduction ability often varies in *in vitro* systems when compared to results from *in vivo* preclinical models in a serotype dependent fashion^18,19^. For instance, AAV8 is very efficient transducing mouse and non-human primate (NHP) livers but it is a poor transducer *in vitro*^20,21^. The leading thought is that AAV attachment factors or receptors in the cell change due to culture conditions.

In this context, we have recently identified a key AAV capsid residue in the variable region I (VR1) (266G) for liver transduction in mice and non-human primates (NHP) using a highly diverse and functional barcoded library of AAV variants based on the ancestral AAV serotype Anc80^22,23^. These findings are in line with likely related observations from various others about the role of VR1 in liver targeting, AAVR binding, and/or epigenetic state of the recombinant genome^24-26^. In our findings, the position 266 within the Anc80 library was associated with strong liver uptake and expression when it was in a Glycine, while an Alanine in that position reduced the liver uptake by over 100-fold. Interestingly, the analogous 266G position in the phylogenetically related AAVrh.74, rh.10, 7, 8 and 9 was conserved, while AAV1, 2 and 3 carried 266A. When we mutated AAV9 to G267A (the aligned position to 266 in Anc80) together with one adjacent change to mimic the Anc80 domain, liver targeting in a mouse model was lost in a significant manner without affecting transduction of e.g. quadriceps. Conversely, when AAV3, generally seen as a poor vector for murine liver, was mutated in this position from A to G (and an additional contextual change), transduction of the liver was enhanced by over 100x. The functional translation of the impact of 266G on human liver transduction remained an open question which we sought to address here.

Specifically, we interrogated relevant human hepatocyte systems both *in vivo* and *in vitro*. To do so, we leveraged the Anc80 library that was extended to include barcoded, naturally occurring and clinically relevant hepatotropic serotypes, such as AAV2, AAV5, AAV8, AAVrh10 and AAVLK03^9,18,26-28^. *In vivo*, we systemically dosed the Anc80 library to FRG mice with a humanized liver with different replacement indexes. *In vitro*, we selected the micropatterned co-culture (MPCC) platform of primary human hepatocytes (PHH) developed by Khetani and Bhatia^29^, where hepatocyte function and phenotype are significantly enhanced for up to 4 weeks by co-culture with mouse embryonic fibroblasts^30^.

By combining these resources, we report that multiplexed NGS analysis of the functional Anc80 library in the human microliver platform yielded results that were reproducible and consistent with the outcomes observed in FRG mice with a high replacement index of human hepatocytes. Importantly, the correspondence between the platforms was dependent on the use of poly(ethylene glycol) (PEG) at the time of *in vitro* hepatocyte transduction, unlocking an AAV entry mechanism usually blocked in hepatocyte cell culture. Our results were also consistent with the AAV transduction pattern observed in the liver of non-human primates, highlighting the specificity of the 266G residue for hepatocyte transduction across species, the potential of the Anc80 library for advancing our understanding on AAV biology, and the predictive potency of the MPCC system for AAV-based human liver gene therapy.

## RESULTS

### Multiplexed AAV library transduction in a xenograft mouse model with a humanized liver

To assess the functionality of the Anc80 A266G liver toggle in the AAV capsid in human hepatocytes as well as its interaction with other functional domains, we tested an extensive, highly diverse library of 2048 vector capsids derived from the ancestral serotype Anc80^21^ in chimeric FRG mice with a humanized liver. This functional library included changes in eleven positions across the Anc80 scaffold sequence where each position has a two amino acid toggle (Figure 1A). In addition, the library included naturally occurring and previously characterized as hepatotropic serotypes AAV2, AAV5, AAV8, AAVrh10 and AAVLK03. For functional readout of gene transfer and transcriptional activity, each of the variants is tagged with a unique barcode that allows for multiplexed DNA and RNA next generation sequencing (NGS)^22^.

**Figure 1.**
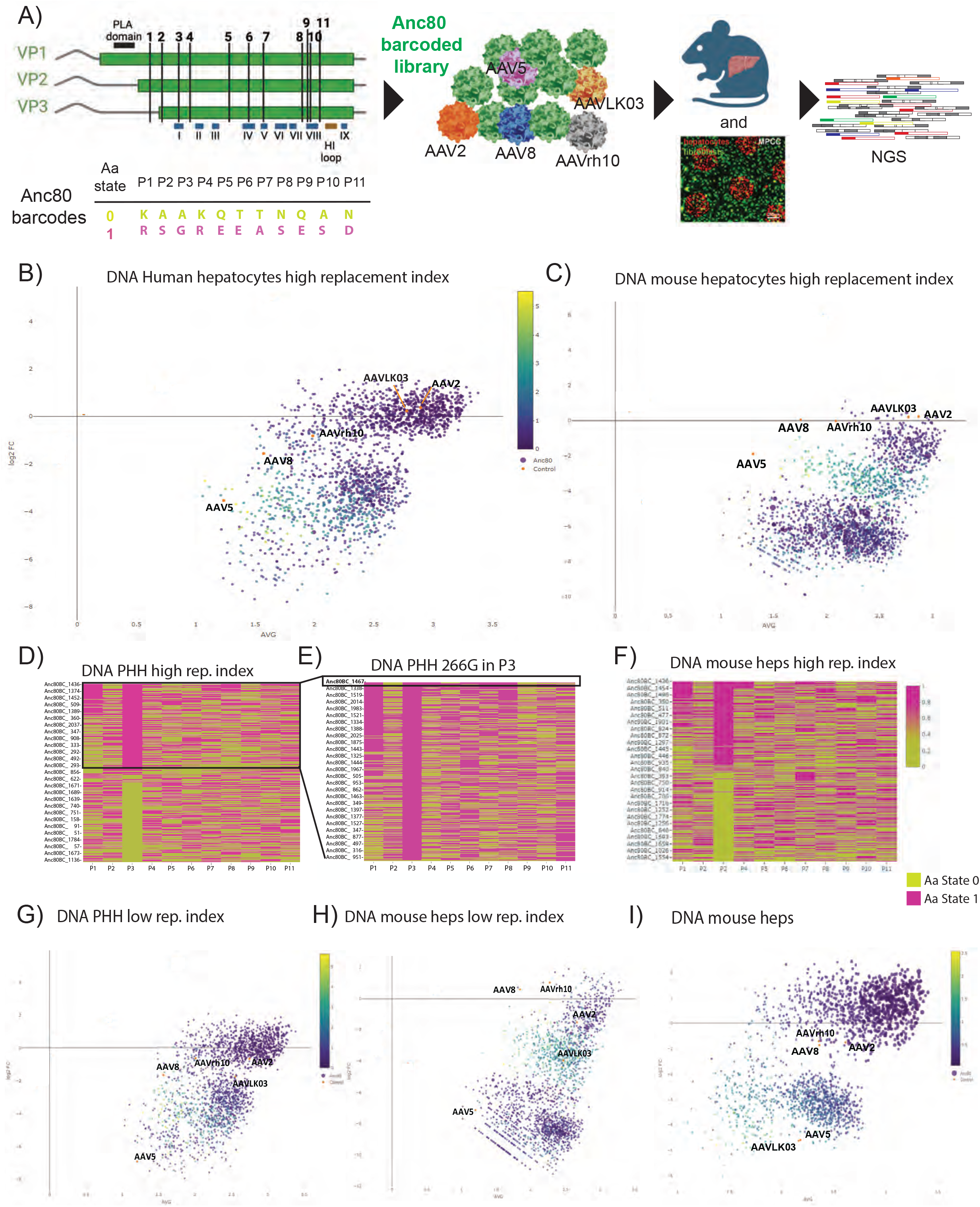
Hepatocyte transduction by the AAV-Anc80 combinatorial library in the humanized liver Fah−/−/Rag2−/−/Il2rg −/− (FRG) mouse model. A) Combinatorial sites of variation in the Anc80 capsid that generated the highly diverse Anc80 library (top), and schematic representation of the experimental design. Bottom left table displays the amino acid toggle for each site of variation and the color code (pink or lime) on the variant ranking. B) MA scatter plot representation of the abundance of DNA barcodes in sorted primary human hepatocytes (PHH) from FRG mice with high repopulation index (n=2). C) MA scatter plot representation of the abundance of DNA barcodes in the mouse hepatocytes from FRG mice with high repopulation index (n=2). Variants are highlighted in a color gradient based on the variance between animals (n=2). High variance (yellow) shows low reproducibility in the barcode readout between animals. Low variance (purple) represents high reproducibility between animals. Barcoded controls are highlighted in orange. D) Ranking of AAV library variants based on DNA barcode enrichment in PHH from FRG mice with high repopulation index. Variants are color coded in lime or pink based on their amino acid state. E) Ranking of AAV library variants with a glycine in position 3 (266G) based on DNA barcode enrichment in D. F) Ranking of AAV library variants based on DNA barcode enrichment in mouse hepatocytes from FRG mice with high PHH repopulation index. G) MA scatter plot representation of the abundance of DNA barcodes in PHH from FRG mice with low repopulation index (n=2). H) MA scatter plot representation of the abundance of DNA barcodes in the mouse hepatocytes from FRG mice with low repopulation index (n=2). I) MA representation of the abundance of DNA library barcodes in the liver of C57BL6/J wild-type mice. Variants are highlighted in a color gradient based on the variance between animals (n=8). Barcoded controls are highlighted in orange.

To control for engraftment bias, we assessed mice with high (>55%) or low (<30%) primary human hepatocyte (PHH) replacement index (Supplementary Table 1). Animals were injected intravenously with 1x10^11^ vector genomes of the highly diverse AAV library per mouse. Four weeks after injection, hepatocytes were harvested from the liver and sorted by species (i.e. human or mouse) before DNA and RNA extraction for NGS analysis. We visualized the abundance of the library members in the PHH (log ratio, y-axis) relative to vector input (mean average, x-axis) using an MA plot (Figure 1B), where enrichment of a given AAV variant is shown as a positive fold change (FC) increment vs the input. The analysis in PHH from mice with high repopulation index revealed an enrichment of a defined population of Anc80 variants, as well as the barcoded controls AAVLK03 and AAV2 (Figure 1B). AAV8, AAV5 or AAVrh10 were not particularly enriched. A lower but similar pattern of enrichment in liver targeting Anc80 capsids (266G) was observed in the sorted mouse hepatocytes (Figure 1C), although no enrichment of the barcoded controls was detected. All variants were ranked and clustered based on their abundance and amino acid combination in the capsid, confirming that the enriched variants in PHH were liver targeting Anc80 capsids with the 266G (Figure 1D-E). Interestingly, the top variants also had an arginine in position 1, located in VP2 (Figure 1E). This entry capsid feature was not human-specific as it was also observed in variants extracted from C57BL6/J mouse livers (Figure 1F). In mice with a lower PHH repopulation index, Anc80 library barcode abundance was like the one in the high replacement index animals (Figure 1G), revealing their preference for entering human cells even though they were less abundant in this context. However, there was a shift in the transduction pattern of the barcoded controls. where AAVLK03 and AAV2 barcodes were reduced in PHH when mice have a low replacement index. Interestingly, AAV5 was strongly de-enriched in the FRG model independently of the replacement index, something also observed by other colleagues^9^. When the mouse hepatocytes were analyzed, we detected an enrichment of AAV8 and AAVrh10 in this species now that more mouse hepatocytes were available for transduction (Figure 1H). Finally, we compared these results to barcode distribution in the non-chimeric C57BL6/J mouse liver, that displayed a similar AAV entry profile to the one in PHH low replacement index but with a strong de-enrichment of AAVLK03 entry (Figure 1I).

### Poly(ethylene glycol) polymer modifies AAV *in vitro* hepatocyte transduction in a predictive fashion recapitulating PHH transduction *in vivo*

To evaluate the 266G liver toggle potency in a reliable PHH *in vitro* system we selected the micropatterned co-culture (MPCC platform), a bioengineered microliver system that has been previously validated for human liver infectious disease applications^30^ or tissue engineering^31^. In the MPCC platform, photolithographic patterning of collagen islands enables adherence of primary hepatocytes of any species origin, which are then subsequently surrounded by mouse embryonic fibroblasts to achieve a controlled ratio of cell-cell interactions^31^. When PHH MPCC were transduced with the highly diverse library (Figure 2A), we could only detect significant enrichment of AAV2 and AAVLK03, something expected due to the high expression of their receptor, HSPGR, in PHH. A modest enrichment of AAV5 barcodes at the DNA level was also observed, but most of the Anc80 library members were not enriched in these conditions nor a specific capsid position in the analyzed variants (Figure 2B).

**Figure 2.**
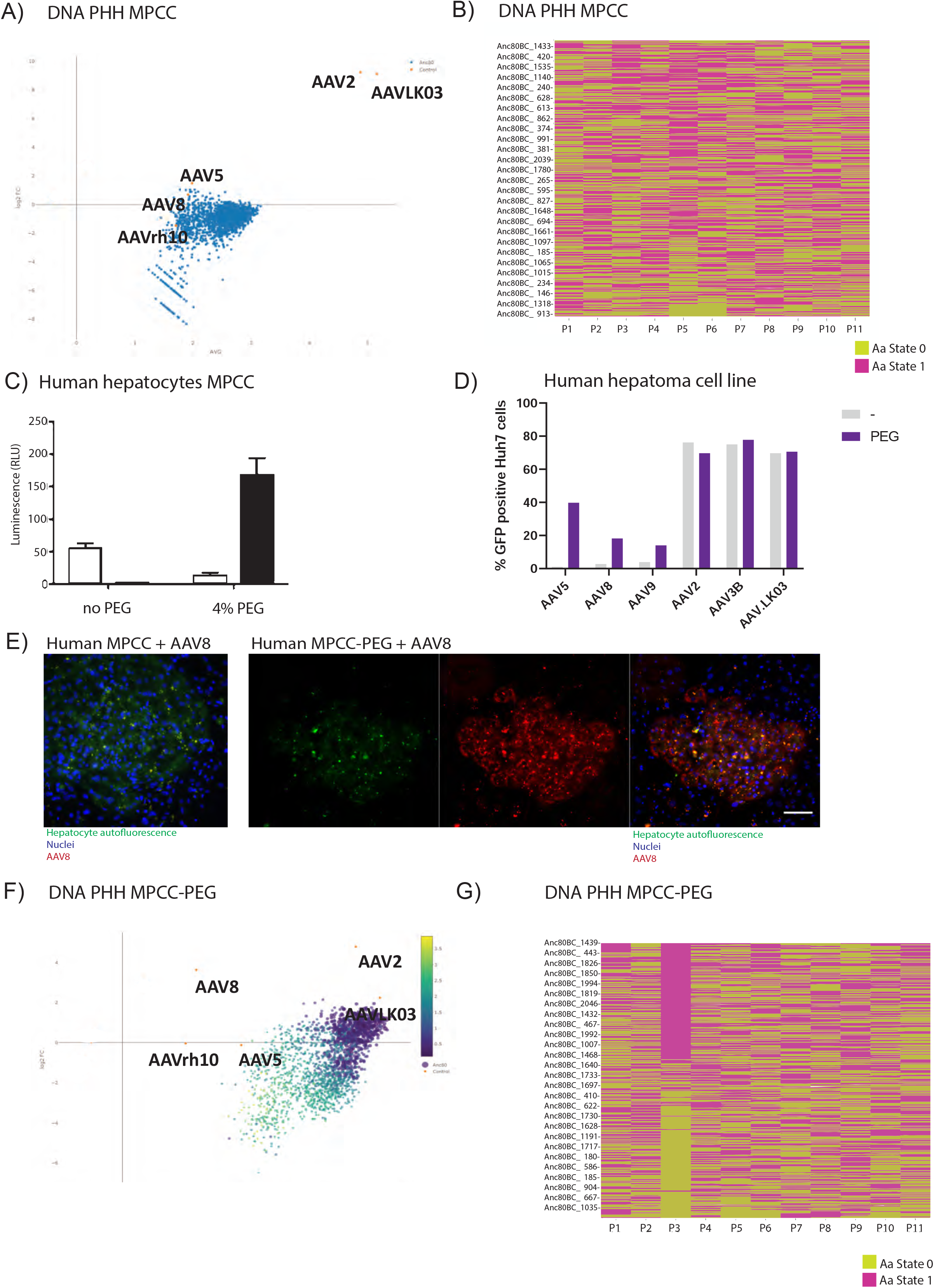
Transduction of primary human hepatocytes in a micropatterned co-culture (MPCC) with a multiplexed AAV-Anc80 combinatorial library. A) MA plot showing DNA barcodes from the AAV combinatorial library in PHH MPCC. On the y-axis, zero indicates no change from the input, with positive and negative values indicating relative enrichment. B) Ranking of AAV library variants based on DNA barcode enrichment in PHH MPCC. Variants are color coded in lime or pink based on their amino acid state. C) PHH were seeded in a MPCC system (96 well plate) and transduced with a MOI of 10^4^ genome copies of AAV2 or AAV8 encoding luciferase, with or without 4% PEG. Data is represented as mean ± sd (n=3 wells) and normalized to AAV2 no PEG condition. RLU, relative luminescence units. D) Human hepatoma Huh7 cells were seeded in a 96 well plate and transduced with a MOI of 10^4^ genome copies of AAV8, AAV2, AAVLK03 and AAV5 encoding EGFP with or without 4% PEG. E) Immunostaining of luciferase in PHH MPCC (left) or MPCC-PEG (right) transduced with AAV8-luciferase. Scale bar = 100um. F) MA plot showing DNA barcodes from the AAV complex library in PHH MPCC with 4% PEG. Variants are highlighted in a color gradient based on the variance between experiments. High variance (yellow) shows low reproducibility in the barcode readout between experiments. Low variance (purple) represents high reproducibility between experiments (n=2, three technical replicates). Barcoded controls are highlighted in orange. (Figure 2A). G) Ranking of AAV library variants based on DNA barcode enrichment in PHH MPCC with + 4% PEG. Variants are color coded in lime or pink based on their amino acid state.

To boost AAV transduction *in vitro*, we investigated biocompatible agents such as polymers that may modify interactions between the virion and the host cell via molecular crowding such as soluble poly(ethylene glycol) (PEG)^32^. First, we tested a 4% (w/v) PEG concentration, which is within the acceptable toxicity limit, combined with AAV2 and AAV8 serotype transduction in MPCC consisting of PHH and mouse fibroblasts (Figure 2C). Hepatocytes were transduced with vectors carrying the firefly luciferase transgene under a constitutively active cytomegalovirus (CMV) promoter. When 4%PEG was added, luciferase activity 48 hours after transduction showed a rank order flip in transduction efficiency, compared to transductions performed without polymer (Figure 2C). To demonstrate robustness and applicability across co-culture formats, we used PHH random co-cultures (RCCs) (Supplemental Figure 1A), hydrogel-laden 3D spheroids of PHH and mouse fibroblasts (Supplemental Figure 1B), or rat hepatocyte MPCC with or without the presence of 4% PEG (Supplemental figures 1B-C). In all these conditions, the addition of PEG resulted in a similar flip of AAV2 and AAV8 rank order. We also tested the AAV+PEG combination in the hepatoma cell line Huh7 and included additional serotypes such as AAV5 and AAVLK03 (Figure 1D). Again, transduction of HSPGR binding serotypes (AAV2 and AAVLK03) was not affected using PEG, whereas AAV8 and AAV5 transduction was enhanced 2-20x times. Luciferase immunofluorescence staining analysis of primary human MPCCs confirmed that AAV8-mediated expression was limited to hepatocytes (Figure 2E).

Next, we assessed the functional relevance of PEG supplementation for AAV entry ability with the highly diverse Anc80 library in the MPCC model. MPCCs consisting of PHH and mouse fibroblasts bearing an inducible apoptotic switch (i.e. inducible caspase-9, which was previously described to be compatible with hepatocyte-fibroblast co-cultures^31^) were transduced with the complex AAV library with and without 4% PEG. 48 hours after transduction, mouse fibroblasts were removed by activating the apoptotic switch to obtain a pure PHH population for DNA and RNA extraction for NGS-based quantification (Fig 2F). By adding PEG, we were able to detect enrichment of more than half of the variant barcodes, including AAV8, together with AAV2 and AAVLK03. We then analyzed the ranking of variants organized by the impact of each toggled position of the viral capsid and ascertained that the PHH transduction change was driven by the 266G toggle in position 3 (Fig 2G).

AAV2 binds to the heparan-sulfate proteoglycan receptor (HSPGR), that is highly expressed in primary hepatocytes *in vivo* and in culture^18^. Canonically, AAV8 is a non-HSPG binding serotype^33^ but given the clear switch in transduction after coating the capsid with PEG, we wondered if PEG was affecting the binding of the capsid to heparan proteoglycans. To address this, the heparin binding capacity of AAV2 and AAV8 with and without 4%PEG was analyzed in a HiTrap Heparin column, as a surrogate of HSPGR binding (Supplemental Figures 1D-G). As expected, AAV2 bound to the heparin column and eluted at a NaCl concentration of 554Mm (Supplemental Figure 1D). When incubated with 4%PEG, the binding of the capsid decreased (Supplemental Figure 1E). These results aligned with the decrease in luciferase expression observed in human hepatocytes when AAV2 is incubated with PEG (Figures 2C-D). We also observed binding of AAV8 to the heparin column (Supplemental Figure 1F), albeit to a lower extend than AAV2 (elution peak at 320mM NaCl concentration). Incubation with 4%PEG completely prevented AAV8 binding to the column (Supplemental Figure 1G-H). These results suggest that the mechanism by which PEG enhances 266G variant hepatocyte transduction *in vitro* is independent of direct HSPGR-mediated entry but instead could be protecting the capsids from strong HSPG attachment and unlocking the interaction between variable region I and their receptor that is observed *in vivo*. Supporting this hypothesis, the top five best performing variants also showed an arginine in position 1 (Supplemental Figure 1I), reproducing the results in Figure 1E.

### Robustness of the MPCC-PEG platform for functional AAV capsid discovery

To understand the functional impact of the 266G residue on transgene expression beyond AAV entry, we investigated RNA barcode distribution in PHH from the *in vivo* (FRG mice with high replacement index) and *in vitro* (MPCC-PEG) models. This is critical because the abundance of AAV genomes in the cell does not always correlate with their transcriptional activity, which is representative of the capsid fitness. Correlation of all the AAV DNA barcodes in the library between these two models was R^2^=0.842 (Figure 3A), whereas correlation score with PHH from FRG with low repopulation index was R^2^=0.752 (Figure 3B). At the transcriptional level, correlation of RNA barcoded transcripts in PHH from the MPCC-PEG and the FRG mice decreased to R^2^=0.736 (Figure 3C). This is because the most potent transducers are usually strong transcribers, but we also observed several variants that had high transcriptional activity with a low hepatocyte transduction profile (Figure 3D). This may be partially explained by the fact that most of the best variants that enter PHH *in vivo* and *in vitro* had a glycine in position 3 and an arginine in position 1 (Figures 1E and Supplemental Figure 1I), but this combination did not display at the RNA level (Figures 3E-F).

**Figure 3.**
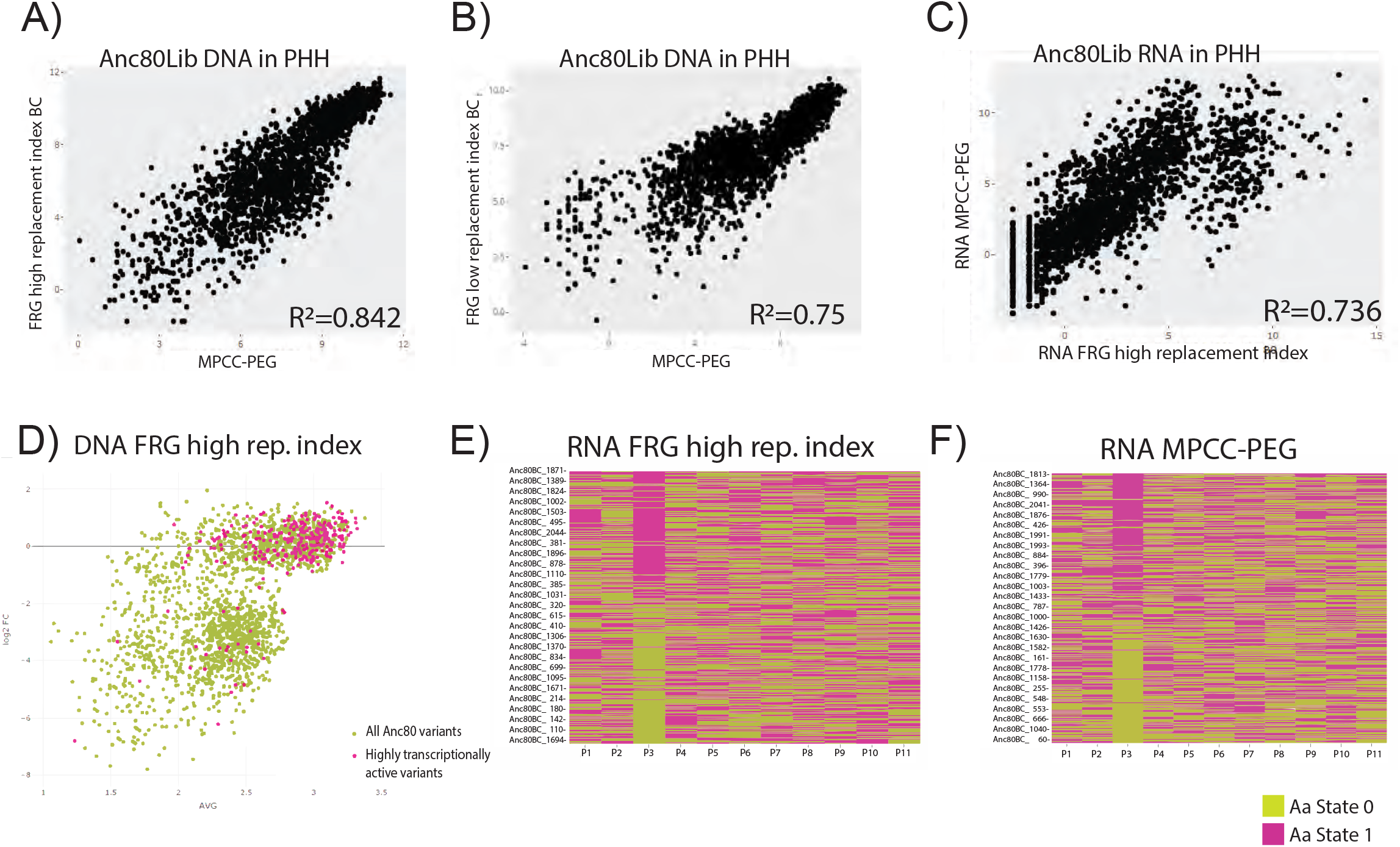
Pearson correlations of DNA barcodes between human hepatocytes from MPCC-PEG and human hepatocytes from the humanized liver of FRG mice with A) high or B) low replacement index. C) MA representation of the abundance of DNA library barcodes in primary human hepatocytes from FRG mice with high repopulation index (lime), where highly transcriptionally active variants have been highlighted in pink. D) Pearson correlation of RNA barcodes between human hepatocytes from MPCC-PEG and human hepatocytes from the humanized liver of FRG mice with high replacement index. E) Ranking of AAV library variants based on RNA barcode enrichment in primary human hepatocytes from FRG mice with high repopulation index. F) Ranking of AAV library variants based on RNA barcode enrichment in PHH MPCC + 4% PEG. Variants are color coded in lime or pink based on their Amino acid state.

Next, we wanted to address specific differences between AAV entry and expression as well as to evaluate the predictive ability of the MPCC-PEG platform for functional AAV capsid discovery. To do so, we selected the best and the worst performing variants at the DNA level in the FRG model with high PHH replacement index, Anc80_1467, liver on, and Anc80_1136, liver-off (Figures 1D-E), and all the naturally occurring AAV barcoded controls. As liver models to compare with, we included PHH from the high replacement index FRG mice, livers from the C57BL6/J mice and livers from two non-human primates (NHP) dosed systemically with the same Anc80 library and sacrificed 4 weeks later^22^. First, all selected variants showed similar PHH transduction efficiency between the FRG humanized mouse model and the MPCC-PEG model (Figure 4A-D), with no significant differences between them but more variability in PHH from the *in vivo* model (Figure 4B). Noteworthy, the Anc80_1136 liver off variant was detected at the DNA level but no transgene expression was found (Figures 4B, D).

**Figure 4.**
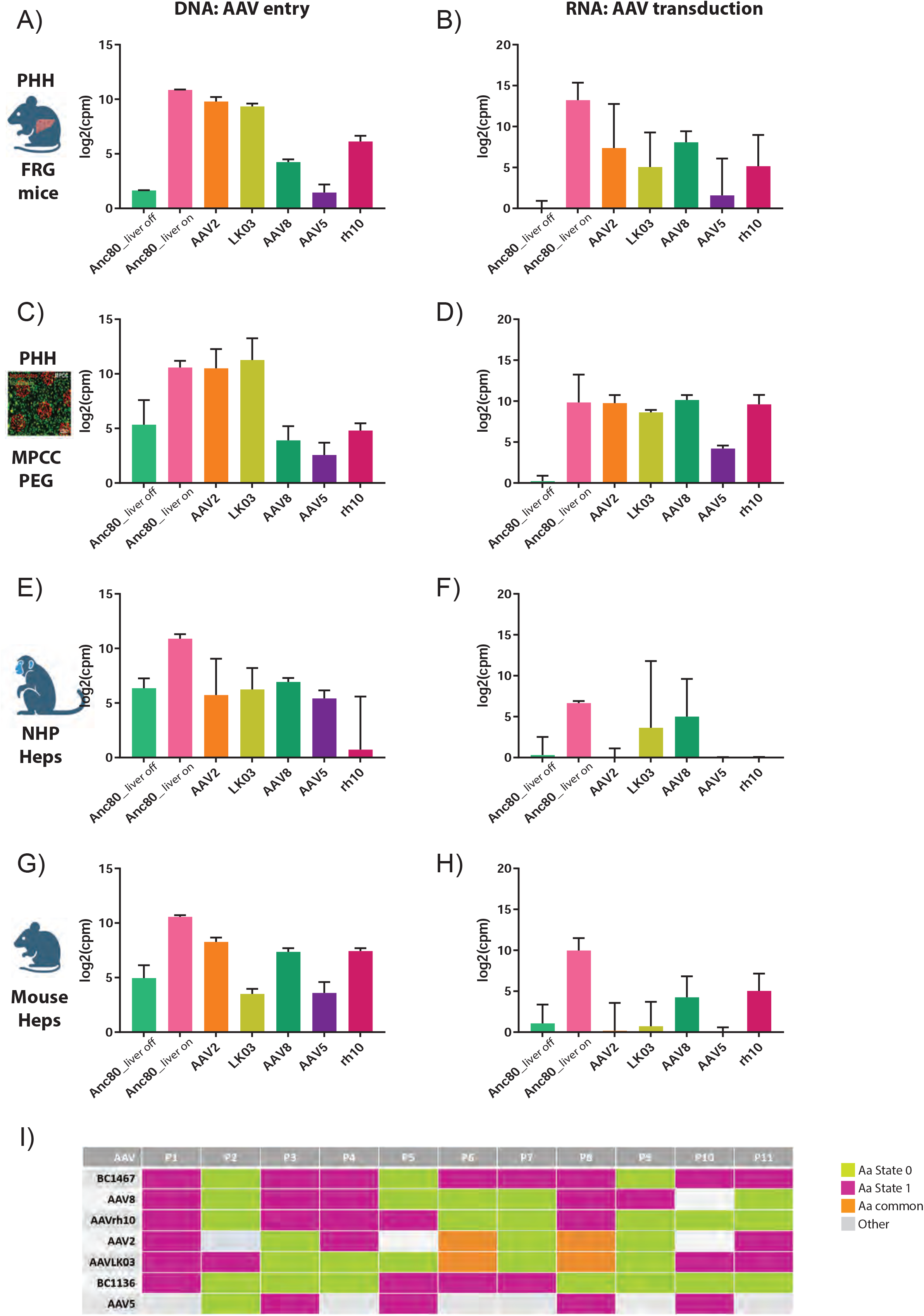
AAV entry versus transcriptional activity from barcoded AAV variants across liver models. Column plot showing the enrichment of DNA barcodes (A) and RNA barcodes (B) of selected AAV variants (log2 of barcodes, counts per million) in human hepatocytes from the humanized liver of FRG mice. C) Enrichment of DNA barcodes and RNA barcodes (D) of selected AAV variants (log2 of barcodes, counts per million) in human hepatocytes from the MPCC+PEG. E) Enrichment of DNA barcodes and RNA barcodes (F) of selected AAV variants (log2 of barcodes, counts per million) in hepatocytes from the liver of non-human primates. G) Enrichment of DNA barcodes and RNA barcodes (H) of selected AAV variants (log2 of barcodes, counts per million) in BL6 mouse livers. I) Comparative table of amino acid states in the Anc80 library variable positions across AAV serotypes of interest. Anc80_BC1467 is the liver on variant and Anc80_BC1136 is the liver off variant. Variants are color coded in lime or pink based on their amino acid state. Orange was used for common amino acids between AAV2 and AAVLK03 in positions 6 and 8. Grey was for other amino acid in that position.

Regarding AAV entry, all selected AAV variants were also detected at the DNA level in NHP and mouse livers, albeit to different proportions as in PHH (Figure 4E, G). However, barcode transcription and functionality changed between these two preclinical models and human hepatocytes (Figure 4F, H). In NHP livers, only the Anc80_1467, liver on variant, AAVLK03 and AAV8 RNA barcodes were clearly detected (Figure 4F), whereas in mice functional transduction was found with the liver on variant, AAV8 and AAVrh10 (Figure 4H). Expression from the liver off variant was consistently abrogated in all preclinical liver models. Comparison of the amino acid state for each AAV variant in the eleven variable positions of the Anc80 library (Figure 4I) identified common residues across liver-on variants, common residues between the liver-off variant and others that did not transduce well NHP and mouse livers, as well as common residues between AAV2 and AAVLK03 in P6 and P8. These results highlight the sensitivity of the combinatorial AAV library as a predictive tool for liver models and AAV biology.

## DISCUSSION

AAV is a leading platform for clinical gene transfer, yet remains with limitations such as sufficient on-target transduction, or conversely excessive biodistribution to off-target tissue, leading to potential liver toxicity^34^. While early studies of basic AAV biology laid the groundwork for therapeutic application, many new challenges have arisen during the translation process. For instance, it was not until recently that the field discovered the biology of different cellular receptors and tropism to different tissue types across naturally occurring AAV serotypes and the key role of the amino-acid sequence in the AAV capsid hypervariable regions^22,23,35^. High-throughput studies of AAV biology requires the development of scalable testbeds with high biological fidelity. However, *in vitro* models to date have generally been unable to capture AAV performance and often reflect inconsistent trends in efficiency when directly comparing preclinical models and human subjects. While AAV libraries are currently mainly used as a high throughput platform for capsid discovery^36^, they are also a powerful tool for unraveling AAV biology. In particular, the use of an AAV library derived from the ancestral Anc80 capsid generated using an *in silico* rationale design led to the identification of a specific position in the AAV genome that directly affect hepatocyte entry in mice and non-human primates (266G)^22^. Our previous work identified the A266G change to be associated with enhanced liver uptake in the mouse and nonhuman primate. In addition, this motif could be introduced by minimal site-directed mutagenesis to convert mouse liver-tropic AAVs to dramatically reduced liver tropism, and vice-versa^22,23^. Intriguingly, these findings added to the importance of VR1, which includes VP1 amino acid 266 in liver targeting and epigenetic state of the recombinant genome in a species dependent manner^25^. This ‘liver toggle’ was proposed as a mechanism to further engineer AAVs to tune the desired tropism to the particular therapeutic application.

To date, it remained unclear whether these observations were restricted to the mouse and nonhuman primate models they were previously studied in. Here, we demonstrate in two human hepatocyte models, *in vivo* and *in vitro*, that the liver toggle effect is also observed qualitatively and quantitatively in these human systems. These findings add to the translational potential of this approach into the clinic.

Furthermore, we have also described for the first time that PEG supplementation at the time of AAV transduction *in vitro* can restore the interactions that AAV variants with a glycine in VP1 amino acid position 266 naturally have with hepatocytes *in vivo.* This is critical because many key interactions between viruses and hepatocytes are lost when these cells are cultured *in vitro*^19,20^. Early on, it was shown that Polybrene (hexadimethrine bromide), a cationic polymer, was able to induce up to a 1000-fold increase in transduction efficiency^37^. Additionally, Gripon et al. showed that supplementing poly(ethylene glycol) (PEG) during infection of hepatitis B virus greatly increased transduction efficiency^32^. Relatedly, it has been shown that functionalization of the AAV capsid with activated PEG can enhance gene transduction, though this method requires chemical modification, leading to attrition of AAV virus stock^38^. In addition, PEG is also used in different AAV production methods for purification^39–41^. However, the addition of a polymer in the context of transduction has never been shown to alter *in vitro* tissue-specific tropism in a predictive fashion. Importantly, the effect of supplementing the inoculation medium with PEG was recapitulated in a range of *in vitro* culture models, from a human hepatoma cell line to a 3D PHH spheroid-laden hydrogels^29-31,36-37^.

Our results showed that PEG-mediated AAV PHH entry *in vitro* was facilitated for those variants with the 266G residue, such as AAV8. On the other hand, PEG supplementation did not affect PHH entry for AAV serotypes that bind the HSPGR, although we observed a decrease in AAV2-mediated transgene expression in several human *in vitro* systems. Therefore, our data suggest that PEG shielding could be blocking potential HSPG-AAV interactions in the hepatocyte extracellular matrix that usually impair their binding to other surface receptors, unlocking their internalization into hepatocytes via AAVR.

Another key feature of our ancestral library is that the barcode was designed to capture both transduction efficacy and transcriptional activity. This additional validation helped identify critical co-dependencies that shape transcriptional activity. For instance, AAVLK03 entered liver cells but did not express the transgene in mouse hepatocytes, as previously described^26^, something we also observed with the Anc80 liver off variant. Recently, it was demonstrated by González-Sandoval and colleagues that the 266 residue in the AAVLK03 capsid was involved in the epigenetic regulation of the episome, and the A266G toggle changed the epigenetic landscape allowing for expression in mouse cells, as in human hepatoma cells^25^. Intriguingly, our data confirmed previous findings with AAVLK03, but in the case of the selected Anc80 liver off variant, barcode expression was abrogated across all tested preclinical liver models. Moreover, AAV5 RNA barcodes were barely expressed in the mouse and NHP livers, suggesting an entry or trafficking disadvantage of this serotype versus those that have a glycine in position 3 (crowding effect), since AAV5 binds N-linked 2,3 sialic acid to enter hepatocytes. Therefore, successful AAV transgene expression is a combination of effective receptor engagement for hepatocyte entry, intracellular trafficking, episome assembly and chromatin accessibility, and residue 266 of the AAV capsid is involved in many of these processes across species, including humans.

The use of the library also allowed us to detect more subtle differences, as the importance of the PHH replacement index to discriminate between good transducers and good transcribers and an intriguing co-selection of 266G in VR1 and 168R in VP2 for efficient hepatocyte entry, but not expression, which further highlights the power of barcode sequencing as a tool to study AAV biology.

From a preclinical model perspective, the FRG mouse is an invaluable tool to bridge data from *in vitro* models and AAV clinical trials, but variability due to differences in the replacement index and cost are bottlenecks for an efficient discovery pipeline. On the other hand, the MPCC platform allows for a more affordable and high throughput scalable system for AAV discovery, and studies can be run for up to four weeks. While co-culture with mouse fibroblasts can complicate downstream analyses that benefit from pure cell populations such as RNAseq and NGS, it was recently shown that the fibroblasts could be genetically modified to carry an inducible apoptotic switch, which could henceforth be activated to remove fibroblasts within 1 hour of dosing and was the system used in our studies^31^. By adding PEG, we were able to reproduce the AAV transduction and transcription pattern observed in a chimeric humanized liver model *in vivo* in the bioengineered MPCC-PHH system. The model accurately reproduced the results from human hepatocyte transduction in FRG mice, showing high correlation in DNA and RNA barcode enrichment. Moreover, we were able to detect new capsid variants within the library that outperformed clinically relevant serotypes in both human preclinical models, as well as variants that were completely PHH detargeted, or may not rely on the liver toggle for human hepatocyte entry but showed high transcriptional activity.

In conclusion, our results showed the potential of a combinatorial AAV library as a tool for model validation and revealed the use of MPCC-PEG as a reliable and predictable human preclinical model for the development of AAV therapeutics. Furthermore, our data suggests, based on findings in two different human models of liver transduction, that the previously described liver toggle may allow for the engineering of vectors to either spare the liver, when the intended target is outside of this tissue, or enhance targeting by rational design.

## MATERIAL AND METHODS

### AAV combinatorial library generation and production

The 2048 barcoded Anc80 capsid library was assembled as described in Zinn et al^22^. To produce high-titer preparations of barcoded Anc80-Lib, the Gene Transfer Vector Core (GTVC) at the Grousbeck Gene Therapy Center of Massachusetts Eye and Ear Infirmary performed quadruple plasmid transfections using polyethylenimine (PEI-Max, Polysciences) in HEK293 cells within ten-layer hyperflasks (Corning). The plasmids were pREP (AAV2 Rep genes under the native p5 promoter), pAAP2 (CMV driving expression of AAP2), the library plasmid (CMV-Barcode-Anc80Capsid), and dF6 (containing necessary Adenoviral accessory genes), in a ratio of 10:10:1:20. Five control barcoded AAVs were spiked into the Anc80 library production keeping the ratio of genome copies per variant in the library. Preparations were tittered by digital-droplet PCR as described in^42^.

### Next generation sequencing

DNA and RNA were extracted from liver cells as detailed below. For barcode sequencing, two step PCR for barcode amplification was performed: barcodes were first amplified by 15 cycles of PCR using primers designed to amplify the barcode and to incorporate a binding site for the subsequent indexing reaction and next barcodes were amplified by 15–35 cycles of PCR, using either primers F1 and R1 (DNA) or F2 and R2 (RNA) for sequencing. To render the amplified barcodes compatible with Illumina sequencing technologies, the purified first-round PCR product was then used as template in a subsequent 8-round indexing reaction using Nextera XT Indexing primers (Illumina Cat No FC-131-1001). PCR products were loaded onto a 2% Agarose gel and were subsequently visualized, extracted, and purified (Zymoclean Gel DNA Recovery Kit, Cat No. D4007). Nucleic acids were quantified by Qubit dsDNA HS Assay Kit (ThermoFisher Cat. No. Q32851). Samples, including the produced multiplexed Anc80 library, were sequenced in a NovaSeq X Plus sequencer at the Broad Institute of Harvard and MIT (Cambridge, MA, USA).

### Barcode analysis and data representation

Data were processed and analyzed using scripts written for this specific purpose as detailed in^22^. Briefly, raw counts were obtained from parallel sequencing lanes together and summed, treating each parallel lane as a technical replicate of sequencing. For each sample within an experiment, the reads are normalized for read depth. To account for variability across biological replicates within an experiment, samples were log-transformed (base 2) and averaged arithmetically. Variance among samples/replicates and fold changes were also calculated as in^22^

### Cell culture

Cryopreserved primary human hepatocytes (Lot ZGF, 33-year-old, male; NON, 35-year-old, female; GEB, 48-year-old, male; and YFA, 10-donor pool aged 6 to 59, mixed gender; all purchased from BioIVT) were thawed in hepatocyte medium without serum and immediately cultured as described below. Primary rat hepatocytes were isolated from 2- to 3-month-old adult female Lewis rats as previously described and seeded in hepatocyte medium without serum^45^. Hepatocyte medium consisted of high-glucose Dulbecco’s modified Eagle’s medium (DMEM) with 4.5 g/L glucose (CellGro) containing 10% (v/v) fetal bovine serum (Gibco), 1% (v/v) ITS (insulin, transferrin, sodium selenite; BD Biosciences), glucagon (70 ng/mL), dexamethasone (0.04 µg/mL), 0.015 M HEPES, and 1% (v/v) penicillin-streptomycin (Invitrogen). Inducible caspase-9 (iCasp9)-expressing 3T3 J2 fibroblasts were prepared from 3T3 J2 murine fibroblasts, a kind gift provided by Howard Green (Harvard Medical School). J2s were lentivirally transduced using a 3rd generation lentiviral system with an iCasp9-IRES-GFP plasmid (Addgene; #15567 pMSCV-F-del Casp9.IRES.GFP; cloned in-house to a lentivirus plasmid backbone with an SFFV promoter), as previously described in depth elsewhere^31^. iCasp9 fibroblasts were maintained in DMEM with 4.5 g/L glucose, 10% bovine serum, and 1% (v/v) penicillin-streptomycin.

### Random and Micro patterned co-culture of primary rat and human hepatocytes

Micropatterned co-cultures (MPCCs) were fabricated as described previously^29^. Briefly, collagen was adsorbed in each well of a 96 well plate (glass bottom; Greiner) and then patterned using an elastomeric polydimethylsiloxane mold and oxygen plasma gas ablation. Primary human hepatocytes were thawed and immediately seeded (70k/well) on collagen islands (500 µm with 1,200 µm center-to-center spacing) in serum-free hepatocyte medium. Alternatively, primary rat hepatocytes were freshly isolated and seeded in a similar manner. Adhered hepatocytes (∼10k/well) were transduced with AAV or a complex AAV library at 1x10^4^ VG/cell prepared in DMEM for 24 hours at 37 C, and supplemented with 4% soluble poly(ethylene glycol) (PEG; Sigma) where indicated. Transduction was performed 0 or 7 days after cell seeding. After transduction, cells were supplied with hepatocyte medium. 24 hours after seeding, inducible iCasp9-expressing J2 fibroblasts were seeded (7k/well) to establish co-culture. 48 hours post-transduction, iCasp9 cells were apoptosed and removed from culture by dosing with B/B homodimerizer (AP20187; TakaraClonTech) at a concentration of 50 nM for 1 hour, as described previously^46^. Hepatocytes were either harvested with 0.25% Trypsin, pelleted, washed with phosphate buffered saline (2-3x), and frozen as pellets for DNA extraction, or lysed with TRIzol® (ThermoFisher) for RNA extraction. For random co-cultures, collagen was adsorbed in each well of a 96 well glass-bottom plate and 25k hepatocytes/well were seeded prior to viral transduction. 24 hours after transduction, 25k fibroblasts/well were co-seeded.

### Spheroid-laden Hydrogel Microliver Culture

Hepatic spheroids were cultured as described previously (^40,42,43^). In brief, cryopreserved human hepatocytes were thawed and immediately plated with fibroblasts in AggreWells (400 micron pyramidal microwells) to enable compaction. Resultant spheroids were encapsulated in fibrin (10 mg/mL bovine fibrinogen, 1.25 U/mL human thrombin; Sigma Aldrich) using 96-well microwell plates as molds. Spheroid-laden hydrogels were cultured in hepatocyte medium supplemented with 10 ug/mL aprotinin, a serine protease inhibitor, to prevent hydrogel degradation.

### Luciferase assay

48h post-transduction, cell culture medium was removed, and cells were lysed in 20 uL per well of 1x Reporter Lysis Buffer (Promega, Cat#E1501), then frozen at -80C. After thaw, ffLuc expression was measured in Relative Light Units/s on a Synergy H1 Hybrid Muli-Mode Microplate reader using 100 uL luciferin buffer [200 mM Tris pH 8, 10 mM MgCl2, 300 uM ATP, 1x Firefly Luciferase signal enhancer (Thermo Cat#16180), and 150ug/mL D-Luciferin].

### Fluorescence microscopy

48h post-transduction, MPCC were fixed with 4% paraformaldehyde for 20 minutes at room temperature, followed by a 3-time wash in PBS. For identification of luciferase-expressing cells, transduced co-cultures were blocked in PBS-BSA 2% for 30 minutes at room temperature and incubated with primary antibody against firefly luciferase (rabbit monoclonal, 1:200; Abcam, ab185924) overnight at 4℃. After PBS washes, cells were incubated for 1 hour at room temperature with goat anti-rabbit IgG (H+L) highly cross-adsorbed secondary antibody, Alexa Fluor 546 (Thermo Fisher, A-11035). Nuclei were stained with Hoechst 33258 (1:5000, Thermo Fisher). Images were captured on a Nikon Eclipse T*i* fluorescence microscope and analyzed with ImageJ.

### DNA and RNA isolation from cell culture and tissue

Total DNA was extracted using the Blood and tissue kit (Qiagen). Total RNA was extracted with TRIzol Reagent (Life Technologies), purified using the PureLink RNA mini kit (Ambion), and treated with RNase-Free DNase (Qiagen) with an on-column DNase treatment.

### Animal experimentation

<u>C57BL6/J</u> animal procedures were performed in accordance with protocols approved by the institutional care and use committees (IACUC) at Schepens Eye Research Institute (SERI). Male C57BL/6 mice (6-8 weeks old) were purchased from Jackson Laboratories and allowed to acclimate to the animal care facilities at SERI for a period of one week and were maintained on a regular diet. <u>FRG</u> (fumarylacetoacetate hydrolase [*Fah^-/-^*] recombination activating 2 [*Rag2^-/-^*] interleukin 2 receptor subunit gamma [*Il2rg^-/-^*]) mice (NOD background) studies were performed at the Children’s Medical Research Institute (CMRI, Sydney Australia). All animal care and experimental procedures were approved by the joint Children’s Medical Research Institute and The Children’s Hospital at Westmead Animal Care and Ethics Committee. CMRI’s established FRG mouse colony was used to breed recipient animals. FRG mice were housed in individually ventilated cages with 2-(2-nitro-4-trifluoro-methylbenzoyl)-1,3-cyclohexanedione (NTBC) supplemented in drinking water (8 mg/ml). FRG mice, 6 to 8 weeks old, were engrafted with human hepatocytes (Lonza Group Ltd., Basel, Switzerland) as described previously (25). hFRG mice were placed on 10% NTBC before transduction with vectors and were maintained on 10% NTBC until harvest. Human serum albumin (HSA) was quantified by ELISA (Bethyl Labs #E80-129). All animals were injected intravenously with 1x10^11^ vector genomes of the Anc80 library per mouse. Four weeks after injection, C57BL6/J livers were collected for nucleic acid extraction and FRG hepatocytes were harvested from the liver by perfusion and sorted by species. Experiments with <u>cynomolgus macaques</u> (Macaca fascicularis) were performed at a contract research organization (DaVinci Biomedical Research, Framingham, MA) in accordance with the institutional IACUCs of both DaVinci and SERI. Animals were pre-screened for antibodies against Anc80L65, AAV1, 2, 5, 8, 9 and rh32.33 and two two-year-old naïve male animals were selected for the study. Animals were injected intravenously with 1.6 ×10^12^ vector genomes/kg of the Anc80 library and sacrificed 28 days after dosing. Tissues were prepared as in the mouse experiments prior to barcode amplification and extraction.

### Sorting of human and mouse hepatocytes

Perfused hepatocytes were labeled with phycoerythrin (PE)-conjugated anti-human-HLA-ABC (clone W6/32, Invitrogen #12-9983-42; 1:20), biotin-conjugated anti-mouse-H-2Kb (clone AF6-88.5, BD Pharmigen #553568; 1:100), and allophycocyanin (APC)-conjugated streptavidin (eBioscience #17-4317-82; 1:500). Flow cytometry was performed in the Flow Cytometry Facility, Westmead Institute for Medical Research (Westmead, NSW, Australia). The data were analyzed using FlowJo 7.6.1 (FlowJo).

### Correlations and statistics

To assess the strength of the linear correlations between the datasets, Pearson correlations were performed using the AAVSeq tool (oculargenomics.meei.harvard.edu/aavseq/), a custom bioinformatics tool for Ancestral AAV library data processing. Statistics to compare individual DNA and RNA variant levels among animal groups were analyzed in Graphpad following a one-way ANOVA test with the Holm-Sídák correction.

## FIGURE LEGENDS

**Supplementary figure 1.** A) Luminescence values at 48h post AAV-Luciferase transduction of a random co-culture of primary human hepatocytes (PHH RCC) with mouse fibroblasts with AAV2 and AAV8 with and without 4%PEG. Bars represent mean ± sd (n=4-6 wells). RLU, relative luminescence units. B) Luminescence values at 48h post AAV-Luciferase transduction of a 3D *in vitro* hepatocyte model: Primary human hepatocytes were aggregated with supporting fibroblasts and transduced with a MOI of 10^4^ genome copies of AAV2 or AAV8, with or without 4% PEG, during compaction. 3D aggregates were embedded in a natural biomaterial matrix and AAV transduction efficiency was assessed 2 days later using the Luciferase Assay System (Promega, E1501). C) Luminescence values at 48h post AAV-Luciferase transduction of a rat hepatocyte micropatterned co-culture (MPCC) with AAV2 and AAV8 with and without 4%PEG. D) AAV2 binding to the HiTrap heparin column measured by ultraviolet absorbance (A280) over time (X-axis). Y-axis represents the linear NaCl gradient in percentage (0-100% = 40-1137 mM NaCl). AAV load into the column is indicated with a blue arrow. Start of the NaCl gradient is indicated with a magenta arrow. AAV elution peak is indicated with a green arrow. E) AAV2 + 4% PEG binding to the HiTrap heparin column. F) AAV8 binding to the HiTrap heparin column. G) AAV8 + 4% PEG binding to the HiTrap heparin column. H) Table of NaCl concentrations at AAV elution peaks. I) Ranking of AAV library variants with a glycine in position 3 (P3) based on DNA barcode enrichment in primary human hepatocytes from the MPCC-PEG system.

**Supplementary table 1.** Summary table of Fah−/−/Rag2−/−/Il2rg −/− mice human repopulation index.

## ACKNOWLEDGMENTS

The authors thank Keval N. Vyas for isolation of primary rat hepatocytes, Eleftherios Michailidis (Rockefeller University) for sharing HBV transduction protocols and Urja Bhatt for supplementary barcode analysis. We also thank Ru Xiao and the Gene Transfer Vector Core (https://www.vdb-lab.org/vector-core/) for manufacturing and providing purified, high-titer adeno-associated viral vectors and libraries used in these studies and Andrea Llanos (CIMA, Universidad de Navarra) for training in FPLC use.

## FUNDING

Funding was provided by the by Giving/Grousbeck to L.H.V. as well as a sponsored research agreement with Lonza Houston (to L.H.V.) and additional support by the British Heart Foundation’s Big Beat Challenge award to CureHeart (award reference no. BBC/F/21/220106). A.X.C. was supported by the National Science Foundation Graduate Research Fellowship (1122374). This work was supported in part by the NIH (R01 EB008396) and the Spanish Ministry of Science and Innovation (RYC2021-033449-I to C.U). Additional funding was provided to the laboratory of S.N.B. by a Koch Institute Support (core) Grant P30-CA14051 from the National Cancer Institute. S.N.B. is a Howard Hughes Medical Institute Investigator.

## DECLARATION OF INTEREST

L.H.V. is a paid advisor to Eli Lilly and Affinia Therapeutics and serves on the Board of Directors of Affinia, Addgene, and Lyora Therapeutics. L.H.V. holds equity in Affinia and Lyora Therapeutics. L.H.V. C.U. and E.Z. are inventors of AncAAV technology licensed to Affinia, Eli Lilly, and/or other biopharmaceutical companies from which they may receive royalties. S.N.B reports compensation for consulting or board membership by Amplifyer Bio, Catalio Capital, Earli Inc., Impilo Therapeutics, Matrisome Bio, Ochre Bio, Port Therapeutics, Ropirio Therapeutics, Satellite Bio, Sunbird Bio, Vertex Pharmaceuticals, and Xilio Therapeutics. L.H.V.’s interests were reviewed and are managed by Mass Eye and Ear and Mass General Brigham in accordance with their conflict-of-interest policies. L.H.V. C.U. and E.Z. are inventors on patent applications relating to the liver toggle technology contained in this publication.

## REFERENCES

1. Zabaleta N, Unzu C, Weber ND, Gonzalez-Aseguinolaza G. Gene therapy for liver diseases - progress and challenges. Nat Rev Gastroenterol Hepatol. 2023 May;20(5):288–305

2. Mendell, J. R. et al. Current Clinical Applications of In Vivo Gene Therapy with AAVs. Mol Ther 29, 464–488 (2021).

3. Costa-Verdera H, Unzu C, Valeri E, Adriouch S, González Aseguinolaza G, Mingozzi F, Kajaste-Rudnitski A. Understanding and Tackling Immune Responses to Adeno-Associated Viral Vectors. Hum Gene Ther. 2023 Sep;34(17-18):836–852.

4. Wang, D., Tai, P. W. L. & Gao, G. Adeno-associated virus vector as a platform for gene therapy delivery. Nat Rev Drug Discov 18, 358–378 (2019).

5. Pupo, A. et al. AAV vectors: The Rubik’s cube of human gene therapy. Mol Ther 30, 3515–3541 (2022).

6. Azuma, H. et al. Robust expansion of human hepatocytes in Fah-/-/Rag2-/-/Il2rg-/- mice. Nat Biotechnol 25, 903–10 (2007).

7. Barzi M, Chen T, Gonzalez TJ, Pankowicz FP, Oh SH, Streff HL, Rosales A, Ma Y, Collias S, Woodfield SE, Diehl AM, Vasudevan SA, Galvan TN, Goss J, Gersbach CA, Bissig-Choisat B, Asokan A, Bissig KD. A humanized mouse model for adeno-associated viral gene therapy. Nat Commun. 2024 Mar 4;15(1):1955.

8. Cabanes-Creus, M. et al. Characterization of the humanized FRG mouse model and development of an AAV-LK03 variant with improved liver lobular biodistribution. Mol Ther Methods Clin Dev 28, 220–237 (2023).

9. Paulk, N. K. et al. Bioengineered AAV Capsids with Combined High Human Liver Transduction In Vivo and Unique Humoral Seroreactivity. Mol Ther 26, 289–303 (2018).

10. Westhaus A, Cabanes-Creus M, Dilworth KL, Zhu E, Salas Gómez D, Navarro RG, et al. Assessment of Pre-Clinical Liver Models Based on Their Ability to Predict the Liver-Tropism of Adeno-Associated Virus Vectors. Hum Gene Ther. 2023 Apr;34(7-8):273–288.

11. Nonnenmacher, M. et al. Rapid evolution of blood-brain-barrier-penetrating AAV capsids by RNA-driven biopanning. Mol Ther Methods Clin Dev 20, 366–378 (2021).

12. Meumann N, Cabanes-Creus M, Ertelt M, Navarro RG, Lucifora J, Yuan Q, Nien-Huber K, et al. Adeno-associated virus serotype 2 capsid variants for improved liver-directed gene therapy. Hepatology. 2023 Mar 1;77(3):802–815.

13. Goertsen D, Flytzanis NC, Goeden N, Chuapoco MR, Cummins A, Chen Y, et al. AAV capsid variants with brain-wide transgene expression and decreased liver targeting after intravenous delivery in mouse and marmoset. Nat Neurosci. 2022 Jan;25(1):106–115.

14. Cabanes-Creus M, Liao SHY, Gale Navarro R, Knight M, Nazareth D, Lau NS, Ly M, Zhu E, Roca-Pinilla R, Bugallo Delgado R, Vicente AF, Baltazar G, Westhaus A, Merjane J, Crawford M, McCaughan GW, Unzu C, González-Aseguinolaza G, Alexander IE, Pulitano C, Lisowski L. Harnessing whole human liver ex situ normothermic perfusion for preclinical AAV vector evaluation. Nat Commun. 2024 Mar 14;15(1):1876.

15. Afonso MB, Marques V, van Mil SWC, Rodrigues CMP. Human liver organoids: From generation to applications. Hepatology. 2024 Jun 1;79(6):1432–1451.

16. Otumala AE, Hellen DJ, Luna CA, Delgado P, Dissanayaka A, Ugwumadu C, Oshinowo O, Islam MM, Shen L, Karpen SJ, Myers DR. Opportunities and considerations for studying liver disease with microphysiological systems on a chip. Lab Chip. 2023 Jun 28;23(13):2877–2898.

17. Shao W, Xu H, Zeng K, Ye M, Pei R, Wang K. Advances in liver organoids: replicating hepatic complexity for toxicity assessment and disease modeling. Stem Cell Res Ther. 2025 Jan 26;16(1):27

18. Petit I, Bernard JS, Faucher Q, Chastagner D, Arnion H, Di Meo F, Védrenne N. Advances in organ-on-chip for drug transporter study: Key insights and hurdles. Drug Metab Dispos. 2026 Jun;54(6):100312.

19. Cabanes-Creus M, Hallwirth CV, Westhaus A, Ng BH, Liao SHY, Zhu E, Navarro RG, Baltazar G, Drouyer M, Scott S, Logan GJ, Santilli G, Bennett A, Ginn SL, McCaughan G, Thrasher AJ, Agbandje-McKenna M, Alexander IE, Lisowski L. Restoring the natural tropism of AAV2 vectors for human liver. Sci Transl Med. 2020 Sep 9;12(560):eaba3312.

20. Wang Q, Firrman J, Wu Z, Pokiniewski KA, Valencia CA, Wang H, Wei H, Zhuang Z, Liu L, Wunder SL, Chin MP, Xu R, Diao Y, Dong B, Xiao W. High-Density Recombinant Adeno-Associated Viral Particles are Competent Vectors for In Vivo Transduction. Hum Gene Ther. 2016 Dec;27(12):971-981.

21. Gao, G. et al. Clades of Adeno-associated viruses are widely disseminated in human tissues. J Virol 78, 6381–8 (2004).

21. Nakai, H. et al. Unrestricted hepatocyte transduction with adeno-associated virus serotype 8 vectors in mice. J Virol 79, 214–24 (2005).

22. Zinn, E. et al. In Silico Reconstruction of the Viral Evolutionary Lineage Yields a Potent Gene Therapy Vector. Cell Rep 12, 1056–68 (2015).

23. Zinn, E. et al. Ancestral library identifies conserved reprogrammable liver motif on AAV capsid. Cell Rep Med 3, 100803 (2022).

24. Cabanes-Creus M, Navarro RG, Liao SHY, Baltazar G, Drouyer M, Zhu E, Scott S, Luong C, Wilson LOW, Alexander IE, Lisowski L. Single amino acid insertion allows functional transduction of murine hepatocytes with human liver tropic AAV capsids. Mol Ther Methods Clin Dev. 2021 Apr 24;21:607–620.

25. Zhang R, Cao L, Cui M, Sun Z, Hu M, Zhang R, Stuart W, Zhao X, Yang Z, Li X, Sun Y, Li S, Ding W, Lou Z, Rao Z. Adeno-associated virus 2 bound to its cellular receptor AAVR. Nat Microbiol. 2019 Apr;4(4):675–682.

26. Gonzalez-Sandoval A, Pekrun K, Tsuji S, Zhang F, Hung KL, Chang HY, Kay MA. The AAV capsid can influence the epigenetic marking of rAAV delivered episomal genomes in a species dependent manner. Nat Commun. 2023 Apr 28;14(1):2448.

27. Lisowski L, Dane AP, Chu K, Zhang Y, Cunningham SC, Wilson EM, Nygaard S, Grompe M, Alexander IE, Kay MA. Selection and evaluation of clinically relevant AAV variants in a xenograft liver model. Nature. 2014 Feb 20;506(7488):382–6. doi: 10.1038/nature12875.

28. Liu S, Razon L, Ritchie O, Sihn CR, Handyside B, Berguig G, Woloszynek J, Zhang L, Batty P, Lillicrap D, Agrawal V, Cortesio C, Gebretsadik K, Akeefe H, Colosi P, Kim B, Bunting S, Fong S. Application of *in*-*vitro*-cultured primary hepatocytes to evaluate species translatability and AAV transduction mechanisms of action. Mol Ther Methods Clin Dev. 2022 May 29;26:61–71.

29. Walkey CJ, Snow KJ, Bulcha J, Cox AR, Martinez AE, Ljungberg MC, Lanza DG, De Giorgi M, Chuecos MA, Alves-Bezerra M, Suarez CF, Hartig SM, Hilsenbeck SG, Hsu CW, Saville E, Gaitan Y, Duryea J Jr, Hannigan S, Dickinson ME, Mirochnitchenko O, Wang D, Lutz CM, Heaney JD, Gao G, Murray SA, Lagor WR. A comprehensive atlas of AAV tropism in the mouse. Mol Ther. 2025 Mar 5;33(3):1282–1299.

30. Khetani, S. R. & Bhatia, S. N. Microscale culture of human liver cells for drug development. Nat Biotechnol 26, 120–6 (2008).

31. Mancio-Silva, L. et al. A single-cell liver atlas of Plasmodium vivax infection. Cell Host Microbe 30, 1048–1060.e5 (2022).

32. Chen, A. X. et al. Controlled Apoptosis of Stromal Cells to Engineer Human Microlivers. Adv Funct Mater 30, (2020).

33. Gripon, P., Diot, C. & Guguen-Guillouzo, C. Reproducible high level infection of cultured adult human hepatocytes by hepatitis B virus: effect of polyethylene glycol on adsorption and penetration. Virology 192, 534–40 (1993).

34. Crosson SM, Bennett A, Fajardo D, Peterson JJ, Zhang H, Li W, Leahy MT, Jennings CK, Boyd RF, Boye SL, Agbandge-McKenna M, Boye SE. Effects of Altering HSPG Binding and Capsid Hydrophilicity on Retinal Transduction by AAV. J Virol. 2021 Apr 26;95(10):e02440–20. doi: 10.1128/JVI.02440-20.

35. Moeini P, Bilbao-Arribas M, Guruceaga E, Torrens-Baile J, Lanz TA, Whiteley LO, Aragón T, Unzu C, González-Aseguinolaza G. DNA damage/p53, innate immune, and unfolded protein responses are activated in primate liver after toxic, high-dose AAV-SMN1 delivery. Mol Ther Adv. 2026 Jan 30;34(1):201682

36. Pillay, S. et al. An essential receptor for adeno-associated virus infection. Nature 530, 108–12 (2016).

37. Wang, D., Tai, P. W. L. & Gao, G. Adeno-associated virus vector as a platform for gene therapy delivery. Nat Rev Drug Discov 18, 358–378 (2019).

38. Chaney, W. G., Howard, D. R., Pollard, J. W., Sallustio, S. & Stanley, P. High-frequency transfection of CHO cells using polybrene. Somat Cell Mol Genet 12, 237–44 (1986).

39. Yao T, Zhou X, Zhang C, Yu X, Tian Z, Zhang L, Zhou D. Site-Specific PEGylated Adeno-Associated Viruses with Increased Serum Stability and Reduced Immunogenicity. Molecules. 2017 Jul 11;22(7):1155.

40. Crosson, S. M., Dib, P., Smith, J. K. & Zolotukhin, S. Helper-free Production of Laboratory Grade AAV and Purification by Iodixanol Density Gradient Centrifugation. Mol Ther Methods Clin Dev 10, 1–7 (2018).

41. Guo, P. et al. A simplified purification method for AAV variant by polyethylene glycol aqueous two-phase partitioning. Bioengineered 4, 103–6 (2013).

42. Arden, E. & Metzger, J. M. Inexpensive, serotype-independent protocol for native and bioengineered recombinant adeno-associated virus purification. J Biol Methods 3, (2016).

43. Stevens, K. R. et al. InVERT molding for scalable control of tissue microarchitecture. Nat Commun 4, 1847 (2013).

44. Stevens, K. R. et al. In situ expansion of engineered human liver tissue in a mouse model of chronic liver disease. Sci Transl Med 9, (2017).

45. Sanmiguel, J., Gao, G. & Vandenberghe, L. H. Quantitative and Digital Droplet-Based AAV Genome Titration. Methods Mol Biol 1950, 51–83 (2019).

46. Seglen, P. O. Preparation of isolated rat liver cells. Methods Cell Biol 13, 29–83 (1976).

47. Davis, H. E., Rosinski, M., Morgan, J. R. & Yarmush, M. L. Charged polymers modulate retrovirus transduction via membrane charge neutralization and virus aggregation. Biophys J 86, 1234–42

