## Supplemental Figure and Table for "Functional evaluation of a natural AAV capsid liver targeting motif in human hepatocytes"

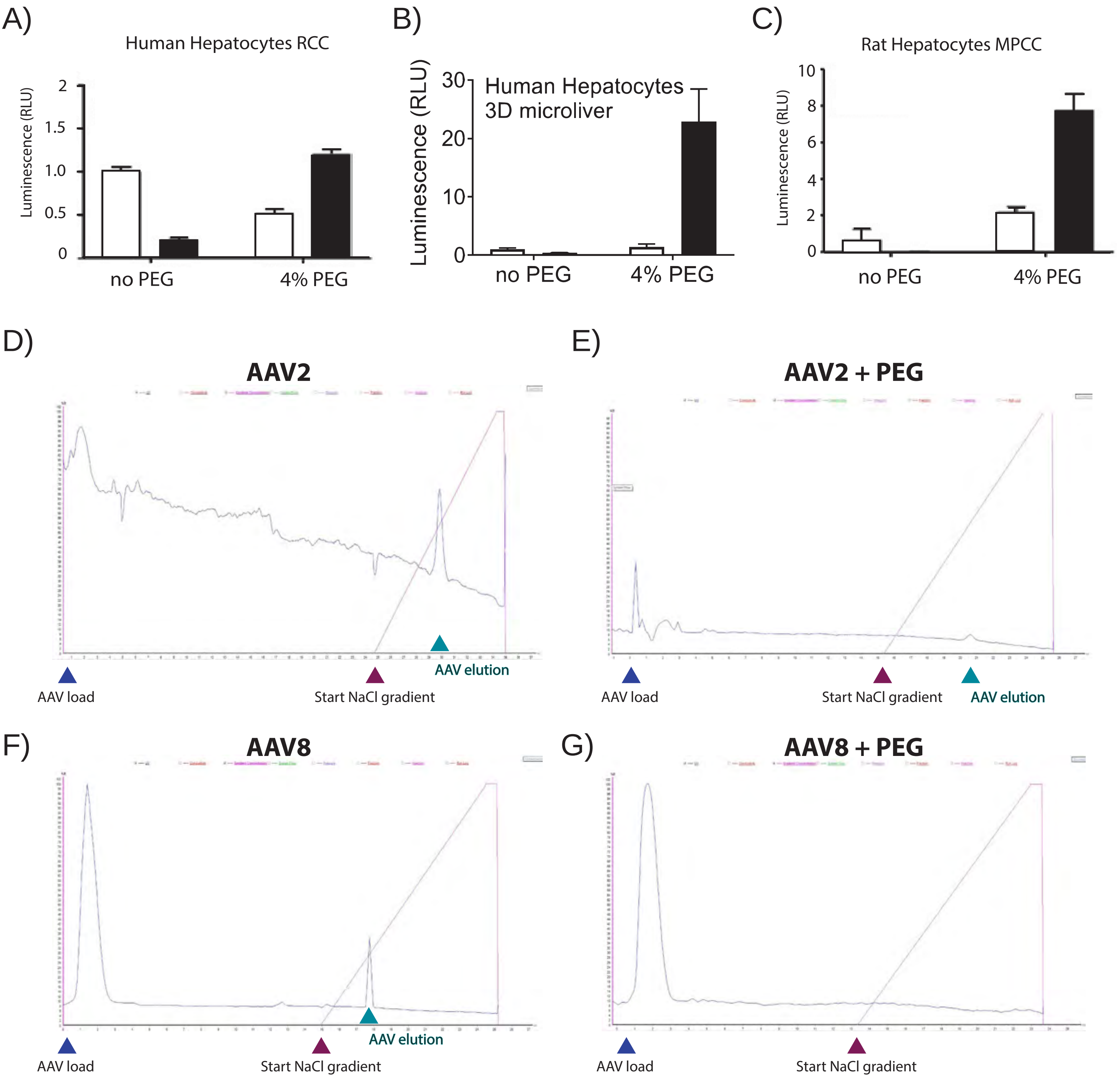


H) **NaCl concentration at AAV elution peak**

| | No PEG | 4% PEG |
|---|---|---|
| **AAV2** | 554mM | 558mM |
| **AAV8** | 328mM | - |

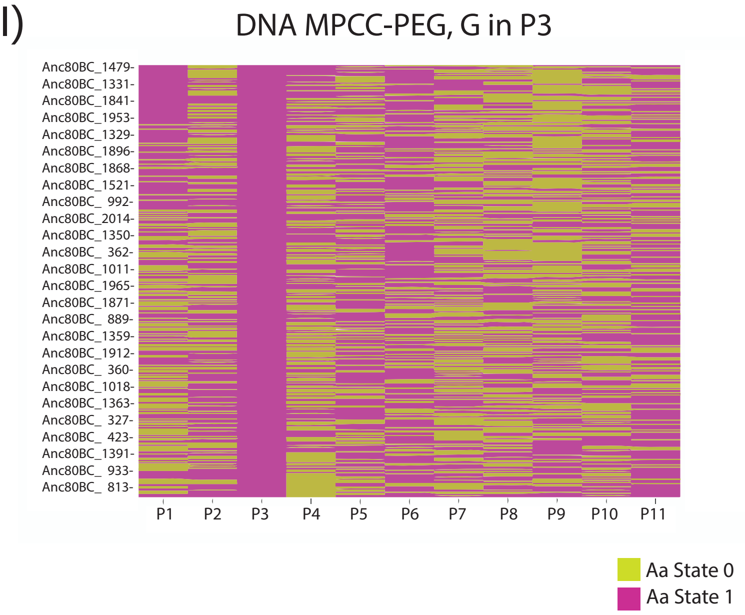


Suppl Figure 1

| mouse ID | human albumin (mg/ml) | % of human hepatocytes by Flow sorting | % of mouse hepatocytes by Flow sorting |
|---|---|---|---|
| 307 | 7.1 | 53 | 38.7 |
| 300 | 7.05 | 65 | 27.84 |
| 368 | 2.14 | 28 | 63 |
| 362 | 2.13 | 36.5 | 50.2 |
